# Emergence of mobile *fosA3-bla_CTX-M-65_* with *mcr-1.1* and *tet*(X4) on conjugative plasmids in avian pathogenic *Escherichia coli* from industrial poultry in Pakistan

**DOI:** 10.64898/2026.11.19.767496

**Authors:** Muhammad Umair, Fariha Fatima, Jordan Mathias, Teresa Iannetelli, Mashkoor Mohsin, Andrew Farlow, Timothy Rutland Walsh

## Abstract

The emergence of plasmid-mediated resistance to critically important antimicrobials poses a growing threat to global health. Here we report the first genomic evidence of plasmid-mediated *fosA3* and *bla*_CTX-M-65_ coexisting with *mcr-1.1* and *tet*(X4) in multidrug-resistant *Escherichia coli* from clinically healthy broiler chickens in Pakistan. Among 1,677 high-quality genomes collected through antimicrobial resistance surveillance in Punjab, Pakistan, three isolates recovered between 2022 and 2024 harboured all four resistance determinants. Hybrid genome assemblies showed that *fosA3* and *bla*_CTX-M-65_ were co-located on *Inc*HI2A plasmids, whereas *mcr-1.1* and *tet*(X4) were carried on separate *Inc*I2 and *Inc*FIA/*Inc*FIC plasmids, all predicted to be conjugative; in one isolate the *tet*(X4) contig did not circularise. The isolates belonged to sequence type 93 and exhibited resistance to fosfomycin, colistin, tigecycline and cefotaxime. Comparative genomic analysis revealed highly conserved *Inc*HI2A plasmid backbones, with an additional integron-associated resistance region in the 2024 isolates, suggesting acquisition of accessory resistance determinants. Resistance to three last-line antimicrobial classes and a canonical virulence profile now coexist on conjugative plasmids in poultry *E. coli*, a combination that favours co-selection and argues for plasmid-resolved surveillance in poultry production.

## Introduction

Food-producing animals (FPAs), including intensive poultry production, account for approximately 73% of global antimicrobial sales, raising concerns about the selection of antimicrobial-resistant bacteria and their potential implications for human health (Van Boeckel et al., 2019). In Pakistan, commercial broiler production relies heavily on medically important antimicrobials, with antimicrobial consumption reaching 462.57 mg/PCU (Umair et al., 2021). The emergence of plasmid-mediated colistin resistance gene *mcr-1.1* in China (Liu et al., 2016) and its subsequent report from Pakistan (Mohsin et al., 2017) brought plasmid-borne colistin resistance to attention as a threat to a last-resort antimicrobial.

Moreover, the emergence of plasmid-mediated tigecycline and fosfomycin resistance genes, *tet*(X4) and *fosA4*, in Pakistan further highlights the growing threat to medically important antimicrobials (Li et al., 2022). Of particular concern is the plasmid-mediated fosfomycin resistance gene *fosA3*, first identified in clinical *Escherichia coli* isolates in Japan (Wachino et al., 2010), and has since been reported in China (Hou et al., 2012) and Brazil (Cunha et al., 2017). Despite its reports from various countries, plasmid-mediated *fosA3* has not previously been reported in Pakistan. Here, we report, the first evidence of the emergence of plasmid-mediated *fosA3* and *bla_CTX-M-65_* in multi-drug resistant (MDR) avian pathogenic *E. coli* (APEC) from healthy broiler chickens in Pakistan, alongside plasmid-borne *mcr-1.1* and *tet*(X4). These findings raise serious One Health concerns and call for enhanced genomic AMR surveillance in commercial broiler production.

### Methodology

As part of an ongoing antimicrobial resistance surveillance study begun in January 2022 on commercial broiler farms in Punjab, Pakistan, cloacal swabs were collected from approximately 3,500 birds, yielding 3,024 *E. coli* isolates. Minimum inhibitory concentrations (MICs) for eight antibiotics were determined using agar dilution for all isolates. Of these, whole-genome sequencing on the Illumina MiSeq platform was performed for 2,021 isolates, yielding 1,677 high-quality genomes. Three isolates carrying multiple clinically important mobile resistance determinants were further characterised using Oxford Nanopore long-read sequencing. Hybrid genome assemblies were generated using Hybracter (Bouras et al., 2024), incorporating Flye, Plassembler and Illumina-based polishing with Polypolish and pypolca/POLCA. Antimicrobial resistance and virulence determinants were identified using AMRFinderPlus (Feldgarden et al., 2021), while plasmid replicons and mobility were predicted using MOB-suite (Robertson and Nash, 2018). Gene-level synteny and circular plasmid comparisons were visualised using clinker and pyCirclize, respectively. Detailed methods are described in Supplementary Material.

## Results

Three *E. coli* isolates carrying *fosA3, bla*_CTX-M-65_, *mcr-1.1* and *tet*(X4) were identified among 1,677 high-quality genomes from a survey of commercial poultry farms in Punjab province, Pakistan. PK-U-0801 was recovered in 2022 from Faisalabad district, whereas PK-U-1553 and PK-U-1591 in 2024 from Khanewal district. All three belonged to sequence type 93 (phylogroup A) and were resistant to apramycin (4–8 mg/L), colistin (8 mg/L), cefotaxime (16–32 mg/L), enrofloxacin (8 mg/L), florfenicol (256 to > 256 mg/L), fosfomycin (MIC 256–512 mg/L), tetracycline (128–256 mg/L), and tigecycline (4–8 mg/L).

Hybrid assembly of Oxford Nanopore and Illumina reads produced closed circular chromosomes of 5,004,689 bp, 4,996,188 bp and 4,996,179 bp, with total assembly lengths of 5.59–5.64 Mb. Within the core-genome alignment of the full collection the three isolates formed a single ST93 cluster separated by 57–177 SNPs, of which 104 lay within a single recombined locus in PK-U-1591. In each isolate the four resistance determinants were distributed across three distinct plasmids rather than co-located on one. *fosA3* and *bla*_CTX-M-65_ were detected on a large *Inc*HI2A plasmid of 265,092–274,296 bp (Figure 1); *mcr-1.1* was the only resistance gene present on an *Inc*I2 plasmid of 60,116–61,043 bp (Supplementary Figure S1); and *tet*(X4) resided on a separate *Inc*FIA/*Inc*FIC plasmid of approximately 94 kb (Supplementary Figure S2). All three replicon types were predicted to be conjugative. In all three isolates *fosA3* and *bla*_CTX-M-65_ lay 4,633 bp apart in the same orientation, flanked by identical genes, and the regions surrounding *mcr-1.1* and *tet*(X4) were likewise conserved between isolates (Figure 2). The *Inc*HI2A plasmid additionally carried resistance determinants against aminoglycosides (*aac(3)-IId*, gentamicin; *aph(3′)-IIa*, kanamycin; and *aph(3′′)-Ib, aph(6)-Ic*, and *aph(6)-Id*, streptomycin), macrolides (*mph(A)* and its regulator *mrx(A)*), sulfonamides (*sul2*), tetracyclines (*tet(A)*) and bleomycin (*ble*), alongside a tellurite-resistance region (*terA*–*terD, terW, terX*, and three copies each of *terY* and *terZ*).

**Figure 1.**
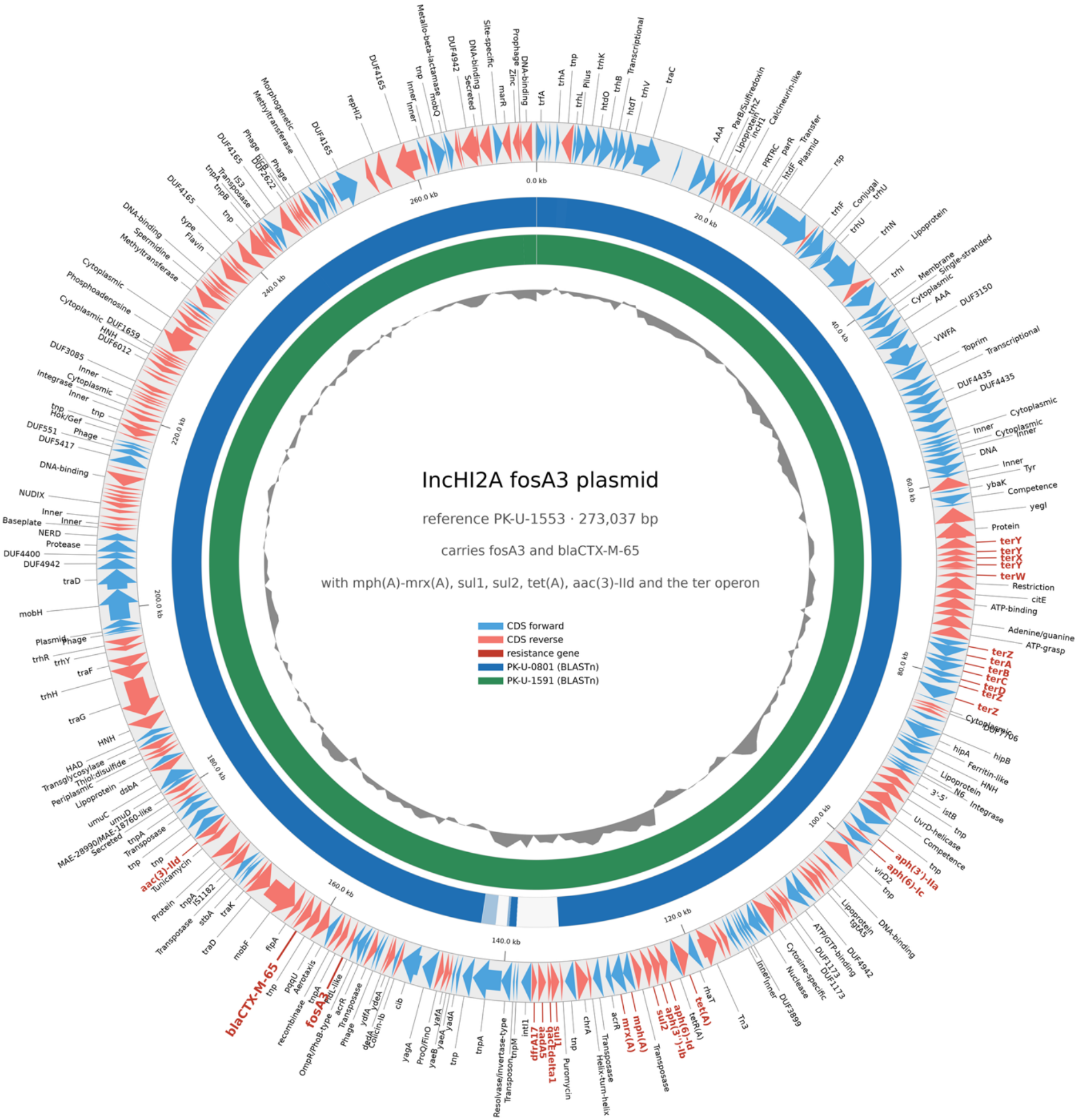
The *Inc*HI2A plasmid carrying *fosA3* and *bla*_CTX-M-65_ is conserved across the three isolates. Circular map of PK-U-1553 (273,037 bp). Outer ring, coding sequences on the forward (light blue) and reverse (red) strands, with resistance genes labelled in dark red; inner rings, BLASTn alignment of PK-U-0801 (blue) and PK-U-1591 (green), where white indicates absence; innermost ring, GC deviation from the mean. *fosA3* and *bla*_CTX-M-65_ lie 4.6 kb apart near 156 kb, within a resistance region that also carries *aac(3)-IId, aph, mph(A)–mrx(A), sul1, sul2, tet(A)* and the ter tellurite operon. The gap in the PK-U-0801 ring at 134 to 143 kb is the 9.3 kb insertion carrying a class 1 integron (intI1, dfrA17–aadA5, qacEΔ1–sul1) within a Tn21/Tn3-family transposon, present in both 2024 isolates.

**Figure 2.**
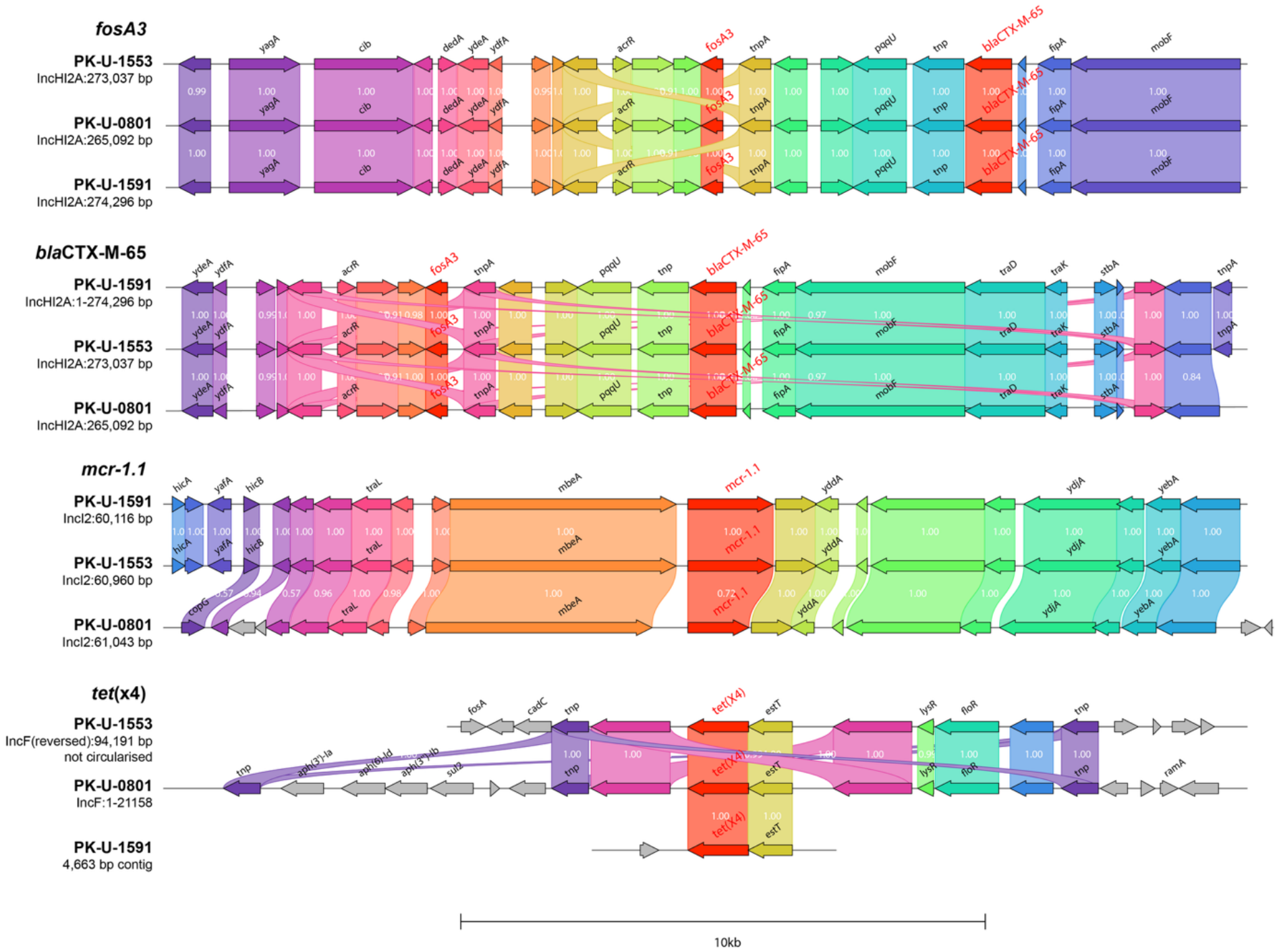
Genetic context of *fosA3, bla*_CTX-M-65_, *mcr-1.1* and *tet*(X4). Genome regions 10 kb either side of each resistance gene, drawn with clinker. Rows are labelled by isolate, replicon type and plasmid size. Arrows are coding sequences; within each panel, genes sharing a colour are homologous and grey arrows have no homologue in the other rows. Colours are assigned separately for each panel and are not comparable between panels. Links give amino-acid identity, and only genes assigned a name by Bakta are labelled. Resistance genes are labelled in red. The PK-U-1553 *tet*(X4) locus is drawn reversed, and the PK-U-1591 window is the whole 4,663 bp contig carrying the gene.

The 265,092 bp *Inc*HI2A plasmid carrying *fosA3* and *bla*_CTX-M-65_ from the 2022 Faisalabad isolate (PK-U-0801) aligned across its entire length with the corresponding plasmids from the two 2024 Khanewal isolates (PK-U-1553 and PK-U-1591), whereas the reciprocal alignment coverage was 97.5% and 97.1%, respectively. This difference was attributable to a 9.3-kb insertion (coordinates 133,760–143,145 in PK-U-1553) comprising a class 1 integron (*intI1*, attI, a *dfrA17–aadA5* cassette array and the 3′ conserved segment *qacEΔ1–sul1–orf5*) embedded within a Tn21/Tn3-family transposon carrying *tnpA, tnpM* and two resolvase genes. These elements were absent from the PK-U-0801 plasmid (Figure 1).

The *mcr-1.1*-carrying plasmids exhibited differences between districts. Plasmids from PK-U-1553 and PK-U-1591 shared 99.977% nucleotide identity, differing by only six single-nucleotide polymorphisms (SNPs) and no insertions or deletions (indels). In contrast, the PK-U-0801 plasmid shared 99.13% nucleotide identity with the Khanewal plasmids, with approximately 400 SNPs, 25 indels and 11–12% unaligned sequence, indicating a closely related but distinct *Inc*I2 plasmid variant (Supplementary Figure S1).

The *tet*(X4)-carrying plasmids of PK-U-0801 and PK-U-1553 shared 99.46% nucleotide identity across 87% of their length (406 SNPs, 38 indels) but carried different accessory resistance genes: *aadA22* (streptomycin and spectinomycin), *aph(3′)-Ia* (kanamycin and neomycin), *aph(3′′)-Ib* and *aph(6)-Id* (streptomycin), floR (phenicols) and *sul2* (sulfonamides) in PK-U-0801, and *bla*_TEM-215_ (β-lactams), *floR* and *fosA4* (fosfomycin) in PK-U-1553. In PK-U-1591 the plasmid was not resolved into a single contig; *tet*(X4) lay on a 4,663 bp contig, and four contigs, two typing as *Inc*FIA/*Inc*FIC, together covered 80% of the PK-U-0801 plasmid at 99.7–100% identity.

A ColV-type *Inc*FIA/*Inc*FIB virulence plasmid, present in all three isolates, carried *cvaB*/*cvaC, iroBCDEN, iss* and *tsh* (Supplementary Figure S3); it was closed at 157,566 and 160,222 bp, the PK-U-1553 contig remaining incomplete at 139,714 bp. With chromosomal aerobactin (*iucABCD, iutA*), *iha, ireA* and *astA*, this constitutes a canonical APEC virulence profile. All three isolates also carried *gyrA* S83L and *parC* S80I, consistent with their enrofloxacin MICs, *bla*EC, and the efflux determinants *acrF, emrE* and *mdtM*. Across all 1,677 genomes, *mcr-1.1* was present in 48.2%, *tet*(X4) in 16.7% and *fosA3* in 3.3% (n = 55); only these three carried all four determinants.

## Discussion

To the best of our knowledge, this study provides the first genomic evidence of plasmid-mediated *fosA3* and *bla*_CTX-M-65_ coexisting with *mcr-1.1* and *tet(X4)* in multidrug-resistant APEC isolates recovered from clinically healthy broilers in Faisalabad (2022) and Khanewal (2024), Pakistan. The high MICs for fosfomycin, colistin, tigecycline and cefotaxime further demonstrate resistance to multiple clinically important antimicrobials.

Importantly, these resistance determinants were distributed across three distinct plasmids predicted to be conjugative: *fosA3* and *bla_CTX-M-65_* on *Inc*HI2A, *mcr-1.1* on *Inc*I2 and *tet(X4)* on *Inc*FIA/*Inc*FIC. Despite the absence of reported fosfomycin use in industrial broiler production in Pakistan (Rehberg et al., 2025), the detection of plasmid-mediated *fosA3* raises concerns about the indirect selection by other antimicrobials. The genetic linkage of *fosA3* with *bla_CTX-M-65_* suggests potential co-selection, whereby *β*-lactam exposure may indirectly favour the persistence and dissemination of fosfomycin resistance, consistent with previous findings in *E. coli* from food-animals (Hou et al., 2013; Jiang et al., 2017; Yang et al., 2014). Furthermore, the conserved *Inc*HI2A plasmid backbone, with an additional integron-associated resistance region in the two 2024 isolates, suggests genetic diversification through the acquisition of additional resistance determinants.

The identification of related ST93 isolates across two districts and sampling years suggests a wider geographical and temporal distribution of this resistant lineage. The 57 to 177 core SNPs separating them are too many for recent direct transmission and point instead to a lineage that has persisted in the production system. Previous studies from Pakistan documented plasmid-mediated *mcr-1.1* and *tet*(X4), while a recent study identified plasmid-borne *fosA4* but chromosomal *fosA3* (Azam et al., 2017; Fatima et al., 2026; Lv et al., 2018). These findings therefore extend the known genetic contexts of fosfomycin resistance in Pakistan. The coexistence of plasmid-borne virulence and resistance determinants in the same APEC isolates indicates that the two are converging in poultry-associated *E. coli*.

Collectively, these findings raise significant One Health concerns regarding the potential dissemination of resistance to last-line antimicrobials and show the need for plasmid-resolved genomic AMR surveillance and strengthened antimicrobial stewardship in commercial poultry production.

## Data Availability

Whole-genome sequence data have been deposited in NCBI under BioProject PRJNA1563645. Raw Illumina and Oxford Nanopore reads for PK-U-0801, PK-U-1553 and PK-U-1591 are available under accessions SRR41826548–SRR41826553, and the three hybrid assemblies under BioSamples SAMN65383221, SAMN65383222 and SAMN65383223. Assembly accessions will be added on release. Supplementary methods, supplementary figures S1 to S3 and the bioinformatics tool versions are provided in the Supplementary Material accompanying this article.

## Acknowledgments

This work forms part of the DPhil thesis of the first author, Muhammad Umair, who gratefully acknowledges funding from the Punjab Educational Endowment Fund, Government of Punjab, Pakistan, for his DPhil studies; the INEOS Oxford Institute for Antimicrobial Research for research funding and for supplementing his stipend to the University of Oxford’s standard rate for one year and nine months; Linacre College for accommodation throughout his DPhil; Linacre College and the University of Oxford for hardship fundings; and the Oxford Pakistan Programme for studentship supports. Muhammad Umair is also thankful to Fariha Fatima, Andrew Farlow, and Timothy Rutland Walsh for their continued support throughout his DPhil. The authors would like to acknowledge the use of the University of Oxford Advanced Research Computing (ARC) facility in carrying out this work http://dx.doi.org/10.5281/zenodo.22558.

## Authors Contribution

MU: conceptualisation, research design, methodology, investigation, data analysis, writing– original draft, writing–review and editing. FF: conceptualisation, methodology, writing–original draft, writing–review and editing. JM: investigation. TL: investigation. MM: writing–review and editing. AF: supervision, writing–review and editing. TRW: funding acquisition, supervision, writing–review and editing.

