## Supplementary Material for "Emergence of mobile *fosA3-bla_CTX-M-65_* with *mcr-1.1* and *tet*(X4) on conjugative plasmids in avian pathogenic *Escherichia coli* from industrial poultry in Pakistan"

**Supplementary Methods**

Cloacal swabs were collected from approximately 3,500 broilers between January 2022 and December 2024 on commercial farms in several districts of Punjab province, Pakistan, as part of a genomic surveillance study of antimicrobial resistance. Each sampled shed housed at least 30,000 birds. Samples were cultured for *Escherichia coli*, and 3,024 isolates were recovered. Minimum inhibitory concentrations of apramycin, colistin, cefotaxime, enrofloxacin, florfenicol, fosfomycin, tetracycline and tigecycline were determined by agar dilution and interpreted according to EUCAST guidelines, with *E. coli* ATCC 25922 as the quality control strain. Of these, 2,021 isolates were whole genome sequenced, of which 1,677 were high-quality genome sequences.

Genomic DNA extracted for PCR was also used for whole-genome sequencing. DNA was quantified on a Qubit fluorometer (Thermo Fisher Scientific) with the High Sensitivity Assay Kit. Short-read libraries were prepared with the Nextera XT kit and sequenced on an Illumina MiSeq using v3 chemistry (2 × 300 bp), giving 31–52× coverage for the three isolates reported here. Long-read libraries were prepared with the SQK-RBK114.96 kit and sequenced on R10.4.1 flow cells (P2 Solo), yielding 124–532 Mb per isolate (22–95× coverage, read N50 11.7–12.2 kb). Reads were base-called in MinKNOW using Dorado in super-accuracy mode (model dna_r10.4.1_e8.2_400bps_sup@v5.2.0). Hybrid genome assemblies were generated using Hybracter v0.14.1 (Bouras et al., 2024), which incorporates long-read quality filtering with Filtlong v0.3.1, de novo assembly with Flye v2.9.6 (Kolmogorov et al., 2019) , replicon reorientation with dnaapler v1.4.0 and small-plasmid recovery with Plassembler v1.9.0 (Bouras et al., 2023). Assemblies were subsequently polished using Illumina short reads with Polypolish v0.7.1 (Wick & Holt, 2022) and pypolca/POLCA v0.5.0 (Zimin and Salzberg, 2020). Medaka polishing was omitted following recommendations for SUP-basecalled reads, and the optimal polishing iteration was selected based on assembly likelihood estimates calculated using ALE, bioconda ale-core 20220503 (Clark et al., 2013).

Antimicrobial resistance and virulence determinants were identified using AMRFinderPlus v4.2.7 with database version 2026-08-07.1 (Feldgarden et al., 2021), while plasmid replicons and mobility were predicted using MOB-suite v3.1.9 (Robertson & Nash, 2018). Plasmid sequences were extracted using SeqKit v2.14.0 (Shen et al., 2016) and annotated using Bakta v1.12.2 database v6.0 (Schwengers et al., 2021). Pairwise plasmid sequence comparisons were performed using dnadiff from MUMmer v4.0.1 (Marçais et al., 2018). Gene-level synteny and sequence similarity were visualised using clinker v0.0.32 (Gilchrist & Chooi, 2021), while circular plasmid comparisons were generated using pyCirclize v1.10.1 with BLASTn 2.17.0+ alignments (Camacho et al., 2009).

### Supplementary Figures

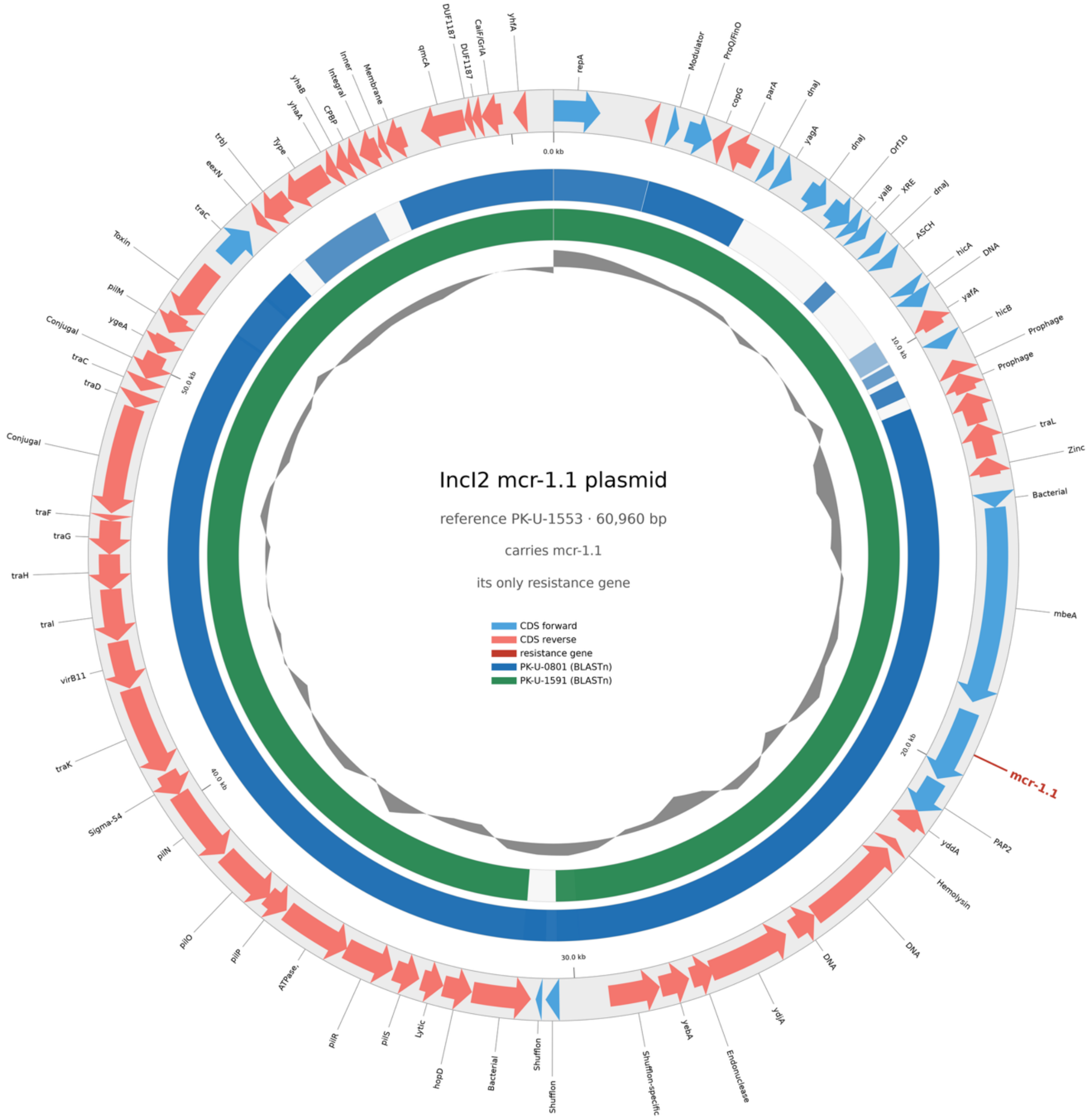


**Supplementary Figure S1.** The *Inc*I2 plasmid carrying *mcr-1.1*, compared across the three isolates.

Circular map of PK-U-1553 (60,960 bp). Outer ring, coding sequences on the forward (light blue) and reverse (red) strands, with *mcr-1.1* labelled in dark red; inner rings, BLASTn alignment of PK-U-0801 (blue) and PK-U-1591 (green), where white indicates absence; innermost ring, GC deviation from the mean. The gaps in the PK-U-0801 ring correspond to the 11 to 12% of sequence unaligned between the 2022 and 2024 plasmids.

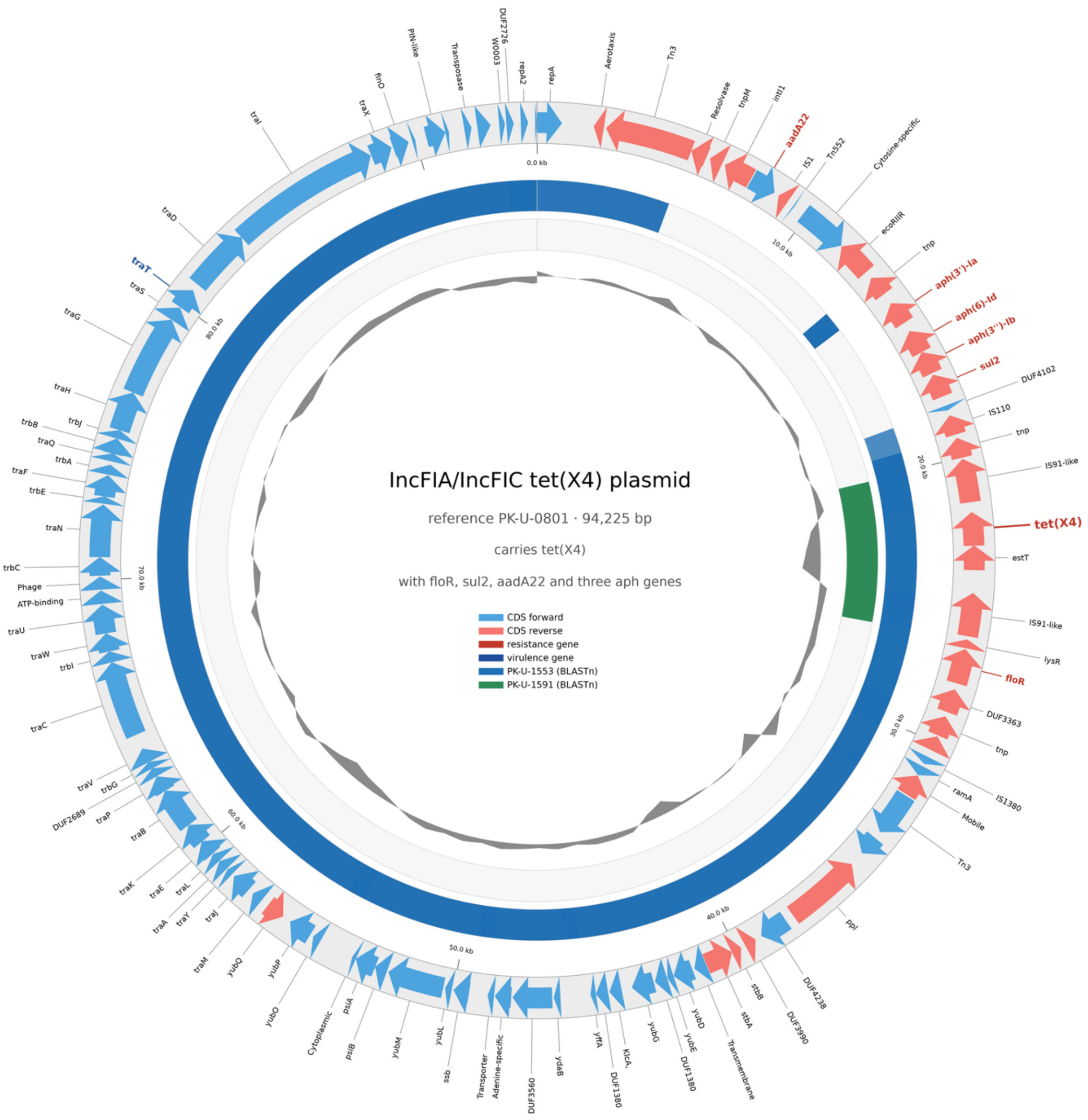


**Supplementary Figure S2.** The *Inc*FIA/*Inc*FIC plasmid carrying *tet*(X4) and accessory resistance genes.

Circular map of PK-U-0801 (94,225 bp). Outer ring, coding sequences on the forward (light blue) and reverse (red) strands, with resistance genes labelled in dark red and the virulence gene traT in dark blue; inner rings, BLASTn alignment of PK-U-1553 (blue) and PK-U-1591 (green), where white indicates absence; innermost ring, GC deviation from the mean. The resistance genes lie between 5 and 30 kb in a region dense with mobile elements (Tn3, Tn552, IS1, IS110, IS91-like, intI1); the break in the PK-U-1553 ring across this region corresponds to the different accessory resistance genes carried by that plasmid. PK-U-1591 contributes only the 4,663 bp contig carrying *tet*(X4), hence the single green arc.

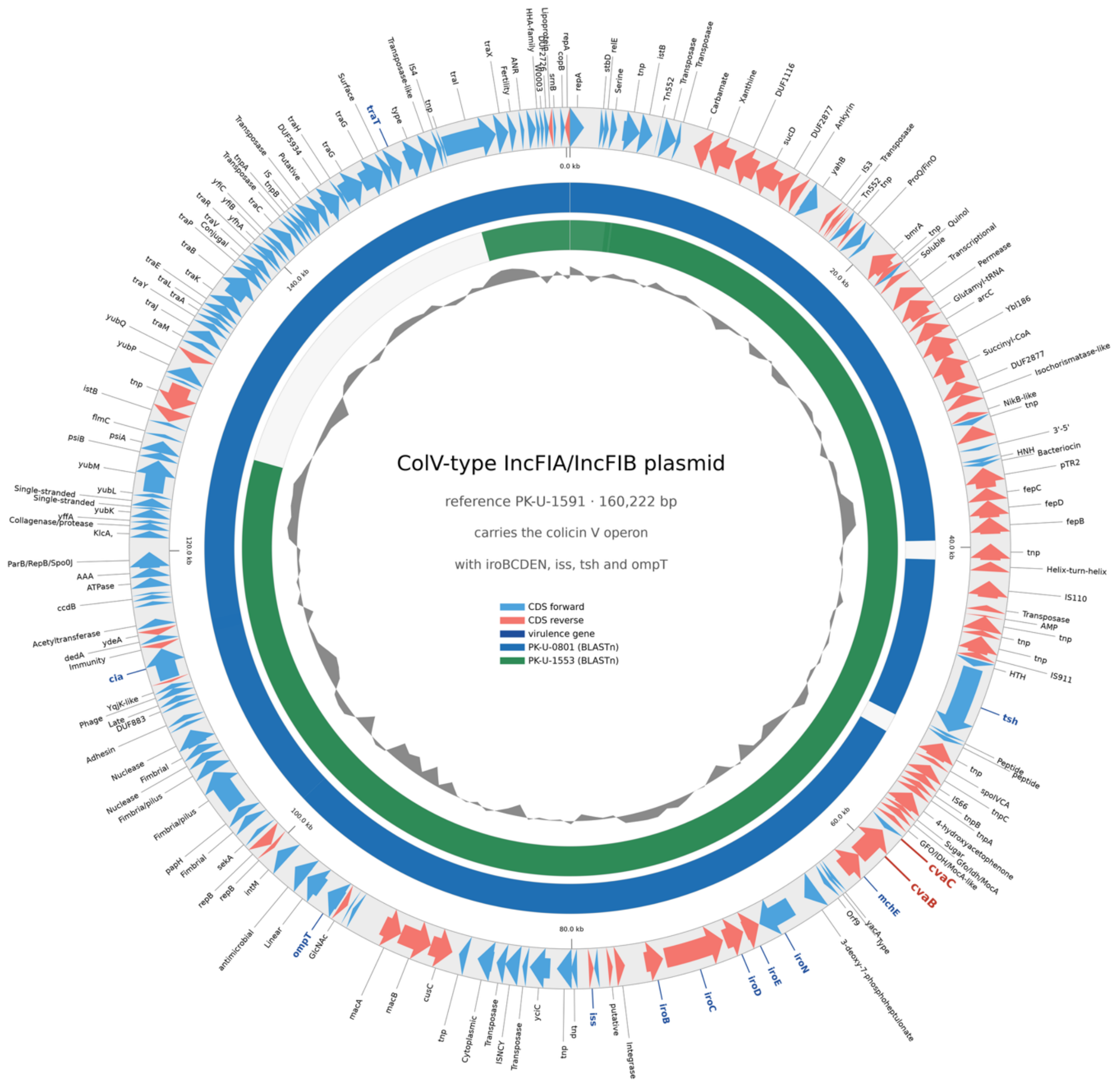


**Supplementary Figure S3.** The ColV-type *Inc*FIA/*Inc*FIB virulence plasmid, present in all three isolates.

Circular map of PK-U-1591 (160,222 bp). Outer ring, coding sequences on the forward (light blue) and reverse (red) strands, with virulence genes labelled in dark blue; inner rings, BLASTn alignment of PK-U-0801 (blue) and PK-U-1553 (green), where white indicates absence; innermost ring, GC deviation from the mean. The virulence determinants lie between 50 and 115 kb (tsh, cvaB/cvaC, mchE, iroBCDEN, iss, ompT, cia), with traT near 155 kb. The PK-U-0801 plasmid aligns across almost the whole reference; the gap in the PK-U-1553 ring between 125 and 140 kb reflects its unclosed contig, not loss of that region.
